# Contrasting evolutionary outcomes in a human life history trait which is heritable and under consistent unbiased directional selection

**DOI:** 10.1101/2025.10.13.682154

**Authors:** Walid Mawass, Emmanuel Milot

## Abstract

Microevolution is well documented in natural populations, yet its persistence as an adaptive process remains debated. Despite widespread directional selection on heritable traits, including life-history traits, evolutionary stasis often prevails, with no detectable microevolutionary response. However, evidence of microevolution in some populations raises a key question: do populations under similar ecological conditions and selective pressures exhibit parallel evolution in the same traits? To address this, we examined age at first reproduction (AFR) in three contemporary human populations considered semi-independent replicates, sharing a genetic, demographic, and historical background, with pedigree data available for ∼7 generations. Across all populations, we found strong directional selection favoring earlier AFR, yet quantitative genetic analyses revealed consistently low heritability (*h*^2^ ≈ 0.11). Only in the Charlevoix population did AFR show a negative genetic correlation with relative fitness, where more than 50% of the standardized phenotypic selection gradient was explained by the genetic selection gradient. Using the Breeder’s Equation and Robertson’s Secondary Theorem of Selection, we predicted an evolutionary response to selection for AFR, which emerged only in Charlevoix. However, neither phenotypic nor breeding values of AFR showed temporal trends, indicating evolutionary stasis. These findings demonstrate that even under consistent directional selection and moderate additive genetic variation, microevolutionary responses may vary across replicate populations. Our results underscore the prevalence of evolutionary stasis, challenging assumptions about the inevitability of microevolution in response to natural selection.

## Introduction

Predicting evolutionary change is a central goal in evolutionary biology (Lässig et al. 2017), with broad implications for understanding population dynamics, adaptation (Hendry et al. 2018), and responses to environmental change (Otto 2018). In human populations, where evolutionary processes intersect with demography (Moorad 2013; Bolund et al. 2015) and public health (Byars and Voskarides 2019; Pavard and Coste 2021), accurate forecasts of trait evolution have tangible consequences for anticipating trends in fertility (Kosova et al. 2010), lifespan (Engelhardt et al. 2019), and population growth (Pelletier et al. 2017). Yet, despite its importance, predicting evolution remains challenging in natural populations. Significant progress has been made largely in experimental systems involving fast-reproducing microorganisms (Neher et al. 2014; Barton et al. 2016), where replication and control are possible.

Contemporary microevolution by natural selection – genetic change in trait distributions over short timescales (Hendry and Kinnison 1999) – is widely thought to occur in nature. This view is supported by experiments demonstrating local adaptation (Barrett et al. 2010; Stuart et al. 2014), by statistical estimates of response to selection from longitudinal phenotypic and pedigree data (Milot et al. 2011; Pigeon et al. 2016; Bonnet et al. 2019), and by the ubiquity of adaptive evolutionary potential based on elevated genetic variance in fitness (Bonnet et al. 2022). However, direct evidence of adaptive evolutionary change in response to environmental variability remains rare (Colautti and Lau 2015; Sauve et al. 2019). One key reason is the difficulty in answering the question of consistency in response to selection both the predictability and repeatability of evolutionary responses. Predictability refers to the alignment between hypothesized responses of heritable traits to selection and their observed temporal trends. Repeatability concerns whether the same phenotypic trait exhibits similar evolutionary outcomes across independent, closely-related populations exposed to similar selective pressures. Despite advances in estimating selection and heritability in the wild, evolutionary responses often lack consistency within and across populations (Grant and Grant 2002; Nosil et al. 2018). In this study, we address the question of consistency by testing the extent of predictability and repeatability of the response to selection.

A major obstacle to predictability is the frequent observation of evolutionary stasis, cases where traits with measurable heritability and directional selection exhibit little or no evolutionary change (Pujol et al. 2018; Wu and Colautti 2022). Understanding the causes of stasis is critical to improving evolutionary predictions. This unpredictability is particularly problematic in modeling human demographic change, where life history traits such as age at first reproduction (AFR) are both heritable and fitness-related. For instance, Pelletier et al. (2017) demonstrated that a substantial portion of the variation in individual contribution to population growth over 108 years in a contemporary human population is attributable to the genetic basis of AFR. In that same population, the mean AFR in married women showed an adaptive response to selection (Milot et al. 2011), suggesting that eco-evolutionary processes influence human population growth. However, if consistent selection on heritable traits fails to yield predictable genetic or phenotypic changes, due to constraints, trade-offs, or stochasticity, then the assumptions underpinning evolutionary demography and eco-evolutionary models may need re-examination.

The consistency problem is further exacerbated by the difficulty in empirically detecting a true genetic response to selection underlying the focal trait. Moreover, selection on a heritable trait does not guarantee a response in breeding values across generations. Phenotypic shifts may occur due to environmental factors, while genetic responses may be masked by opposing environmental trends, a phenomenon known as cryptic evolution (Hadfield et al. 2011). Conversely, evolutionary stasis may be caused either from biological constraints, such as genetic correlations (Teplitsky et al. 2014), pervasive stabilizing selection (Hansen and David 2004), phenotypic plasticity (Nussey et al. 2007), indirect genetic effects (Kruuk et al. 2015), and fluctuating selection (Siepielski, DiBattista, and Carlson 2009; Siepielski et al. 2013; but see Morrissey and Hadfield 2012).

Quantitative genetic (QG) models, such as the animal model, provide tools for estimating, with adequate longitudinal data, heritabilities, selection gradients, and temporal trends in breeding values (Kruuk 2004; Wilson et al. 2010). These models allow researchers to distinguish, to a certain extent, between phenotypic and genetic change, and to formally test whether observed trait variation is consistent with microevolution. Moreover, approaches based on Robertson’s Secondary Theorem of Selection (STS) offer a robust way to estimate the genetic response to selection, compared to the classic Breeder’s Equation (Lush 1937; Lynch and Walsh 1998), by measuring the additive genetic covariance between a trait and relative fitness (Robertson 1966; 1968), without requiring full knowledge of the causal pathway from trait to fitness (Morrissey et al. 2012).

In humans, large-scale genealogical and demographic records provide unique opportunities for such analyses. Empirical evidence from genomic and phenotypic studies confirms ongoing selection on heritable traits like reproductive timing (Byars et al. 2010; Tropf et al. 2015; Sanjak et al. 2017; Stern et al. 2021), demonstrating the evolutionary potential of complex traits in contemporary human populations.

The historical French-Canadian (FC) population presents a valuable system for testing the consistency of evolutionary responses. This population underwent rapid demographic expansion via colonization and founder events during the preindustrial period (Moreau et al. 2011; Peischl et al. 2018; Anderson-Trocmé et al. 2023). Life history theory predicts that such conditions should favor early reproduction, especially in high-fertility, growing populations (Cole 1954; Stearns 1992; Roff 1993). In the Île aux Coudres (IAC) population, this prediction has been supported empirically: directional selection for earlier AFR was accompanied by a downward shift in the population mean, which was partly due to a genetic response to selection (Milot et al. 2011). However, this population is endemic to a small island and its population scale is small compared to the overall French-Canadian population to which it belongs. Furthermore, Mawass, Mayer, and Milot (2022) showed that at the scale of the island, rapid environmental fluctuations can affect the response to selection in AFR as indicated by G×Es underlying the additive covariance between AFR and relative fitness.

In this study, we extend this analysis to a larger population scale by focusing on three subregional FC populations, Bois-Francs, Charlevoix (to which IAC belongs to, historically), and Côte-de-Beaupré, that share genetic, ecological, and demographic histories. These “natural replicates” allow us to ask a fundamental question: When selection acts consistently on a heritable trait, do evolutionary outcomes follow a predictable pattern? To answer this, we use QG models to: i) estimate the heritability of AFR among married women, ii) quantify selection and predict evolutionary change, and iii) detect any underlying shift in breeding values across generations.

Surprisingly, our results reveal diverse evolutionary outcomes across the three populations despite consistent selection for earlier AFR which is demonstrably heritable. In one population, the trait evolved as predicted; in another, it remained unchanged; in the third, it shifted in the opposite direction. However, in all three populations, no significant temporal trend in the average breeding values in AFR was detectable. These contrasting results highlight the challenges of forecasting evolutionary change, even under seemingly ideal conditions, and underscore the importance of integrating genetic data with demographic models to better understand the limits and scope of evolutionary prediction.

## Materials and methods

### Study system

The historical French-Canadian (FC) population is uniquely characterized by detailed parish and civil records documenting marriages, baptisms, and burials. Since the 1970s, these records have been systematically compiled and linked into two comprehensive databases: BALSAC (*http://balsac.uqac.ca/*; Bouchard et al. (1989); Vézina and Bournival (2020)) and the Ancient Québec Population Register (AQPR; Dillon et al. (2018)). These datasets provide extensive pedigrees, enabling the reconstruction of individual and family life histories for descendants of the French settlers who established New France in 1608. We leveraged this rich resource to conduct QG analyses.

The demographic expansion of the FC population followed a series of founder events and subsequent growth, leading to sub-regional populations well-suited for QG studies. We selected three preindustrial populations with high-quality pedigrees and detailed life history records (Mawass and Milot 2025). The dataset for each population was curated to include individuals born and married in the following regions before 1833: Charlevoix, located ∼100 km northeast of Québec City on the north bank of the St. Lawrence River, Côte-de-Beaupré, bordered by Québec City to the west and Charlevoix to the east, and Bois-Francs (Centre-du-Québec), situated on the south shore of the St. Lawrence, approximately midway between Québec City and Montréal. These regions were among the earliest to receive French settlers migrating outward from the main settlement of Québec City.

Our analysis focused on *age at first reproduction* (AFR) among women. To ensure consistency, we included only women who married once and had at least one recorded reproductive event. An additional filter based on the age of last reproduction was applied to establish an upper reproductive limit, coinciding with natural fertility cessation around menopause (typically ∼45 years; Gold (2011)). These criteria align with those used in previous studies (Milot et al. 2011; Pelletier et al. 2017; Mawass et al. 2022). Table 1 presents descriptive statistics for AFR across the three datasets.

**Table 1.** Average values of married women’s life-history traits and standard deviations.

| Trait | Charlevoix<br>(N=2226) | Côte-de-Beaupré<br>(N=3181) | Bois-Francs<br>(N=4120) |
| --- | --- | --- | --- |
| Age at first birth (y) | 23.4 ± 4.6 | 24.4 ± 5.0 | 23.9 ± 4.9 |
| Age at last birth (y) | 36.1 ± 6.3 | 35.2 ± 7.3 | 33.1 ± 7.5 |
| Infant mortality (offspring death<br>between 0-1 y) | 0.9 ± 1.4 | 0.3 ± 0.7 | 0.2 ± 0.5 |
| Fertility (completed family size) | 7.2 ± 3.7 | 6.3 ± 3.9 | 5.03 ± 3.6 |
| Reproductive value at birth | 1.9 ± 1.0 | 3.8 ± 3.0 | 3.1 ± 3.12 |

While these populations span comparable time periods, specific criteria were used to define study periods: the lower bound corresponds to the earliest birth year in the filtered datasets and the upper bound reflects the last year of birth with comprehensive life history data available. In Charlevoix, the time period ends before significant outmigration to the Saguenay-Lac-St-Jean region, which depleted the adult population. Accordingly, we defined the last birth year as that of women born and married in Charlevoix before this demographic shift. The resulting study periods are as follows: Bois-Francs (1688–1833), Charlevoix (1683–1816) and Côte-de-Beaupré (1661– 1832). This temporal delineation was essential for accurately calculating relative fitness. Records predating the study periods—covering marriages, baptisms, and burials—were fully incorporated to reconstruct associated pedigrees.

### Fitness measurements

We measured relative fitness, *w*, for each known married woman as her reproductive value at birth (Fisher 1930; Lenski and Service 1982). Fitness was calculated relative to vital rates in the studied population:

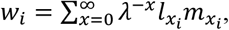

where *w*_*i*_ is the fitness of woman *i*, 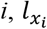 is the survival to age *x*, 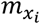 is the age-specific fertility at age *x*, and *λ* is the intrinsic rate of population growth (Moorad 2014). Due the low data density at each discrete year, calculating *λ* separately for annual cohorts would be burdened by high imprecision (earlier years comprised less than <10 married individuals). Therefore, we calculated *λ* for 20-year cohorts. Note that mean fitness in each population is > 1 (Table 1) due to post-estimation filtering to include only women who had at least one reproductive event.

### Quantitative model fitting

All model fitting was conducted within a Bayesian framework to avoid the complications of mixing Bayesian and frequentist approaches, which can lead to inconsistencies when interpreting parameter estimates. In Bayesian analysis, the posterior distribution represents the probability of causes (i.e., the likelihood that a parameter assumes specific values), in contrast to the frequentist approach, which focuses on the probability of effects (i.e., data) given the causes. Models were fitted using the MCMCglmm package (Hadfield 2010) for R (v4.2.1; R Development Core Team 2022). We assumed a Gaussian error structure for all models, and priors were set to default in MCMCglmm, using a Gaussian distribution for the fixed effects and an inverse-gamma prior for the random effects.

### Estimating phenotypic selection gradient

We investigated the relationship between AFR and relative fitness at the phenotypic level using mixed-effect linear regression models. Both the response variable (AFR) and covariates were standardized (mean = 0, standard deviation = 1) to obtain standardized selection coefficients (Lande and Arnold 1983). We fitted both linear (*β*) and quadratic (*γ*) effects to estimate linear and non-linear selection gradients. The equation of the model is as follows:

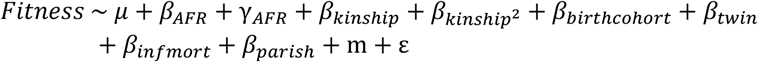

The fixed effects included: whether a woman had at least one pair of twins, birth cohort, birth parish, and the number of infants who died before age 1. Previous studies showed a moderate level of inbreeding in the Charlevoix population (∼0.007; Gérard Bouchard, Braekeleer, and SOREP (1991)), and in île aux Coudres, inbreeding was non-linearly related to infant mortality (Boisvert and Mayer 1994). Therefore, we included the quadratic effect of the kinship coefficient in the model. A random effect for maternal identity was also included to account for the familial environment shared by siblings, given the monogamous nature of the couples.

### Bivariate quantitative genetic model

We calculate trait heritability, estimate selection based on Robertson’s covariance, and detect potential genetic responses to selection in AFR, using the “animal model” (Henderson 1975; 1986), a linear mixed-effects model that incorporates the pairwise genetic relatedness matrix from the pedigree, using all levels of relatives (siblings, grandparents, aunts/uncles, etc.). The animal model also allows for separating the genetic component of phenotypic change from non-genetic components (Wilson et al. 2010).

We first fitted bivariate models with AFR and fitness as response variables to decompose their phenotypic variance into additive genetic, environmental, and residual components, and to estimate the genetic covariances among these traits for each dataset. The model for this analysis was:

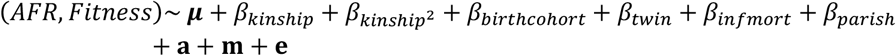

where ***z*** is the response trait vector, ***μ*** is a vector of means, **a** and **m** represent the additive genetic effects and shared familial effects, respectively, while **e** is the residual error matrix and *β* is a vector of partial regression coefficients for each fixed predictor. Additionally, we included as fixed effects whether a woman gave birth to at least one pair of twins and the number of her infants who died before age 1 (infant mortality count), as well as the focal individual’s inbreeding coefficient as a covariate (quadratic effect). The woman’s birth cohort and birth parish were entered as covariates to control for temporal trends or spatial effects that may be due to unmeasured factors.

Narrow-sense heritability (*h*^2^) was estimated as the ratio of the additive genetic variance 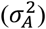 to the total variance in the model 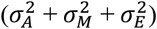. However, as narrow-sense heritability does not always effectively compare the capacity for selection response across populations, we also calculated the coefficient of additive variation 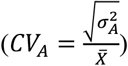, which measures the evolvability of a trait relative to its mean (Houle 1992).

We ran two types of analyses: 1) with mean-centered response variables to obtain variance estimates on the observed scale (for calculating the coefficient of variation using the non-standardized mean) and 2) standardized (mean = 0, standard deviation = 1) to obtain standardized estimates of (co)variances. All response variables were treated as Gaussian (see Figures S2 and S3 for the raw distributions of AFR and fitness in the three datasets). To minimize autocorrelation in the MCMC chain, all models were run for 3,000,000 iterations after a burn-in of 500,000 iterations, sampling every 3,000 iterations to obtain the posterior distributions (1,000 iterations kept). We used an inverse-Wishart distribution for the prior on the variance-covariance matrices (De Villemereuil et al. 2016).

### Bias in phenotypic selection gradients

To assess potential biases in the phenotypic selection gradients, we calculated an “environmental bias” metric, which quantifies the difference between the environmental selection gradient (*β*_*E*_) and the genetic selection gradient (*β*_*G*_). The environmental selection gradient is the regression slope of environmental deviations for fitness on environmental deviations for age at first reproduction (AFR), while the genetic selection gradient is the regression slope of breeding values for fitness on breeding values for AFR (Rausher 1992; Stinchcombe et al. 2014). This approach recognizes that phenotypic covariance between fitness and a focal trait can arise purely from environmental factors (e.g., nutritional status) without any causal influence of the trait on fitness (Price et al. 1988). In such cases, the phenotypic selection gradient may be biased, even though the trait might be heritable, suggesting no real selection on the underlying genetic values. Consequently, no response to selection would be expected, as the environmental covariance does not reflect selection on the breeding values.

The environmental bias in selection was denoted as *Δβ* = *β*_*E*_ − *β*_*G*_, where the posterior distribution of this difference was extracted from the Bayesian models. To calculate *β*_*G*_ and *β*_*E*_ for AFR, we used the following equations:

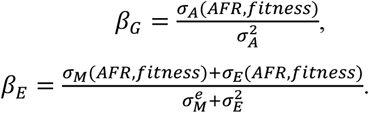

where *σ*_*A*_, *σ*_*M*_, and *σ*_*E*_ represent estimates of the additive genetic covariance, maternal (familial) covariance, and environmental covariance components between AFR and fitness, respectively, with the corresponding variances estimates for each dependent variable.

### Interpretation of posterior distributions

The 95% Highest Posterior Density (HPD) interval provides a measure of uncertainty about the true parameter value. It is important to note that the HPD interval is not to be confused with a frequentist confidence interval. A wider HPD interval suggests greater uncertainty or lower precision in the estimate of the parameter. In addition to the HPD interval, we utilized two Bayesian indices to draw inferences from the posterior distribution about the existence and significance of genetic or environmental effects on the traits: the *region of practical equivalence* (ROPE)-based index and the *probability of direction* (pd).

The ROPE-based index (specifically the ROPE-95%) indicates the proportion of the 95% HPD interval that lies within the ROPE. The ROPE defines a range of values that are considered negligible or practically irrelevant to the biological question at hand (Kruschke 2014). This index is a Bayesian alternative to the frequentist concept of “power” and helps to assess whether the data supports the null hypothesis of no effect. The ROPE-based index is sensitive to sample size and provides insight into the practical significance of the effect (Makowski, Ben-Shachar, Chen, et al. 2019).

*pd* is the Bayesian equivalent of the frequentist p-value and reflects the likelihood that an effect exists in a particular direction. This index ranges from 50% to 100%, with higher values suggesting that an effect is likely present, and lower values indicating greater uncertainty about the effect’s existence. The pd is a direct measure of the presence of a meaningful effect in the data. Both indices were calculated using the bayestestR package in R (Makowski, Ben-Shachar, and Lüdecke 2019), offering robust tools for assessing the significance and direction of genetic and environmental influences on the studied traits.

### Microevolutionary response to selection

In this study, we employed both Breeder’s Equation (BE) and Robertson’s Secondary Theorem of Natural Selection (STS) (or Robertson’s covariance) to predict the expected microevolutionary change in AFR between generations, expressed in terms of phenotypic standard deviation. The distinction between BE and STS lies in their assumptions about environmental factors. BE assumes that environmental influences do not affect the covariance between fitness and the trait. If this assumption is violated, BE may fail, while STS can still be valid, as it directly predicts the response to selection based on the relationship between breeding values (BVs) for the trait and relative fitness (Morrissey et al. 2012).

BE describes the expected evolutionary response in a trait from one generation to the next, given the narrow-sense heritability of the trait and the selection pressure acting on it. The equation is as follows:

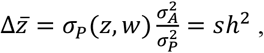

where 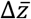 is the predicted change in trait mean phenotype after one generation, *s* represents the phenotypic selection differential and *h*^2^ is the narrow-sense heritability amounting to the proportion of phenotypic variance explained by the additive genetic variance.

STS states that the evolutionary change is equal to the additive genetic covariance of a trait with relative fitness:

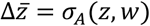

To validate any detected temporal trend in BVs as a response to selection, we used a Bayesian method that calculates the likelihood of observing the slope of the trend in BVs under random genetic drift. Genetic drift can introduce changes in breeding values (BVs) over time that are not driven by selection, so it is crucial to isolate this effect when testing for evolutionary responses. The method involves using the posterior estimates of 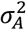 from the sampled MCMC iterations to generate random breeding values along the pedigree, simulating a scenario where no selection is acting on the trait. These random breeding values are then regressed against the birth cohort, and the slope of this regression is compared to the slope obtained from the regression of the model estimated BVs.

## Results

### Phenotypic selection

The results of the linear mixed models, which includes the standardized linear (*β*) and quadratic (*γ*) selection gradients for AFR, revealed a strong linear selection on AFR younger values across all three populations (Table S1). Thus, a younger AFR is associated with higher values of fitness in all the analyzed populations. The standardized selection gradient on AFR was strongest for Charlevoix with *β* = −0.64 [95% HPD: −0.68 – −0.60], then for Côte-de-Beaupré *β* = −0.34 [−0.37 – −0.30], and for Bois-Francs *β* = −0.25 [−0.29 – −0.22] (Figure 1 and Table S1). Interestingly, we also detected a slight quadratic selection gradient in the Charlevoix population *γ* = 0.07 [0.05 – 0.08], but this parameter was negligible for the two other populations (Table S1).

**Figure 1.**
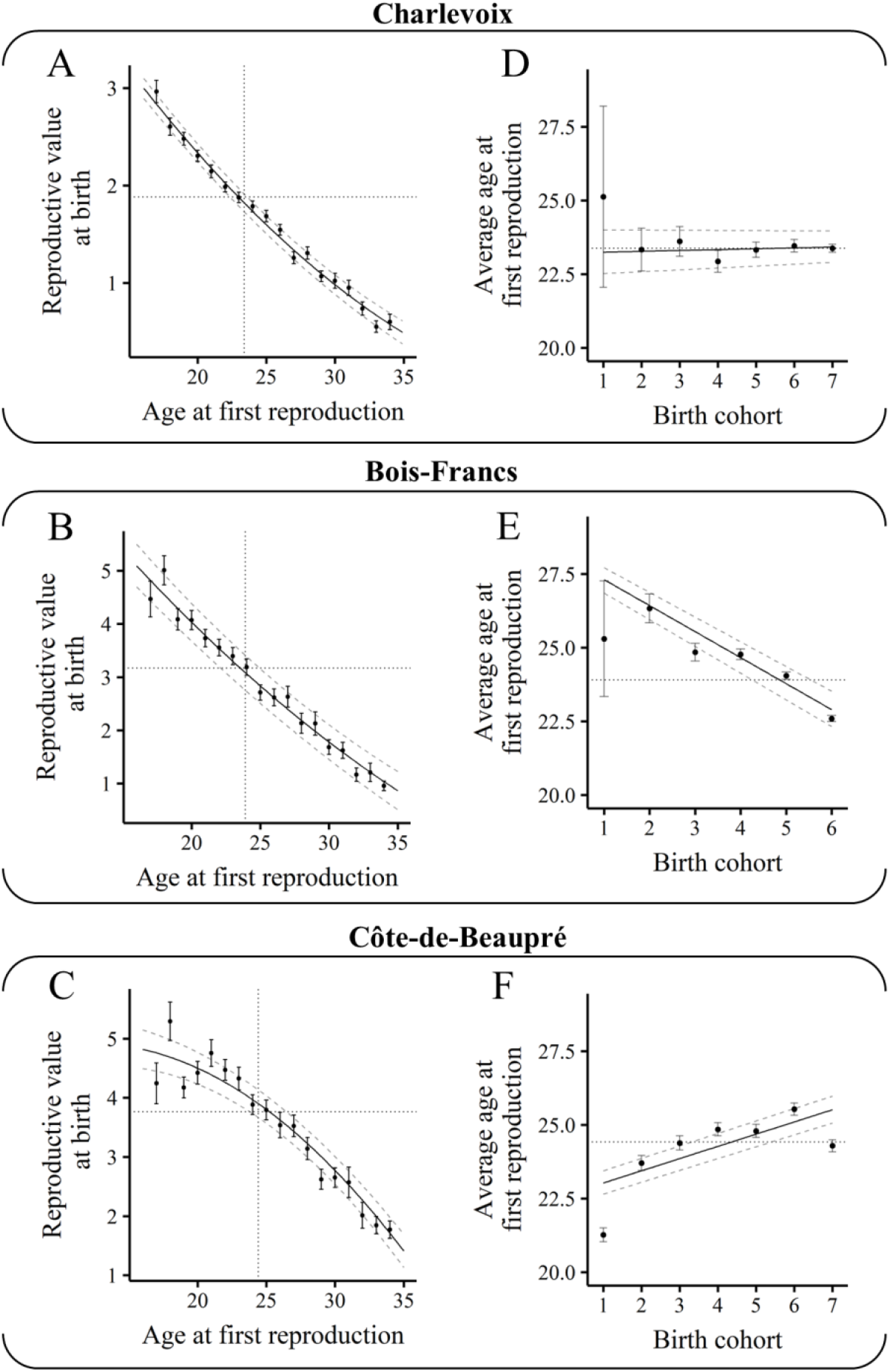
Consistent and strong selection for earlier AFR across all populations, but temporal trend in average AFR differ. All regression lines are based on predictions made using the best-fit linear mixed-effect model. The dashed lines represent the 95% HPD interval of the regression line. Dots represent the observed average values. Vertical and horizontal dotted lines represent population-specific averages.

### Quantitative genetic parameters

In this section, we detail the results of the bivariate animal model for each studied population. We report the results based on the standardized models to allow for comparison at the same scale. The bivariate model shows moderate heritability in both AFR and relative fitness (see Table 2.2 and 2.3, and Figure 2). Across all three populations, the shared familial (maternal) environment was found to be an important contributor of variance to the phenotypic variation in AFR and relative fitness. The coefficient of additive genetic variation was modest for AFR in Charlevoix (CV_A_ = 0.15 [0.06 – 0.26]) and very low in Bois-Francs (CV_A_ = 0.03 [0.02 – 0.06]) and Côte-de-Beaupré (CV_A_ = 0.04 [0.02 – 0.06]). Surprisingly, the coefficient of additive genetic variation was very low for relative fitness in Charlevoix (CV_A_ = 0.05 [0.03 – 0.08]), while it was higher in Bois-Francs (CV_A_ = 0.24 [0.15 – 0.35]) and Côte-de-Beaupré (CV_A_ = 0.17 [0.10 – 0.23]).

**Table 2.**
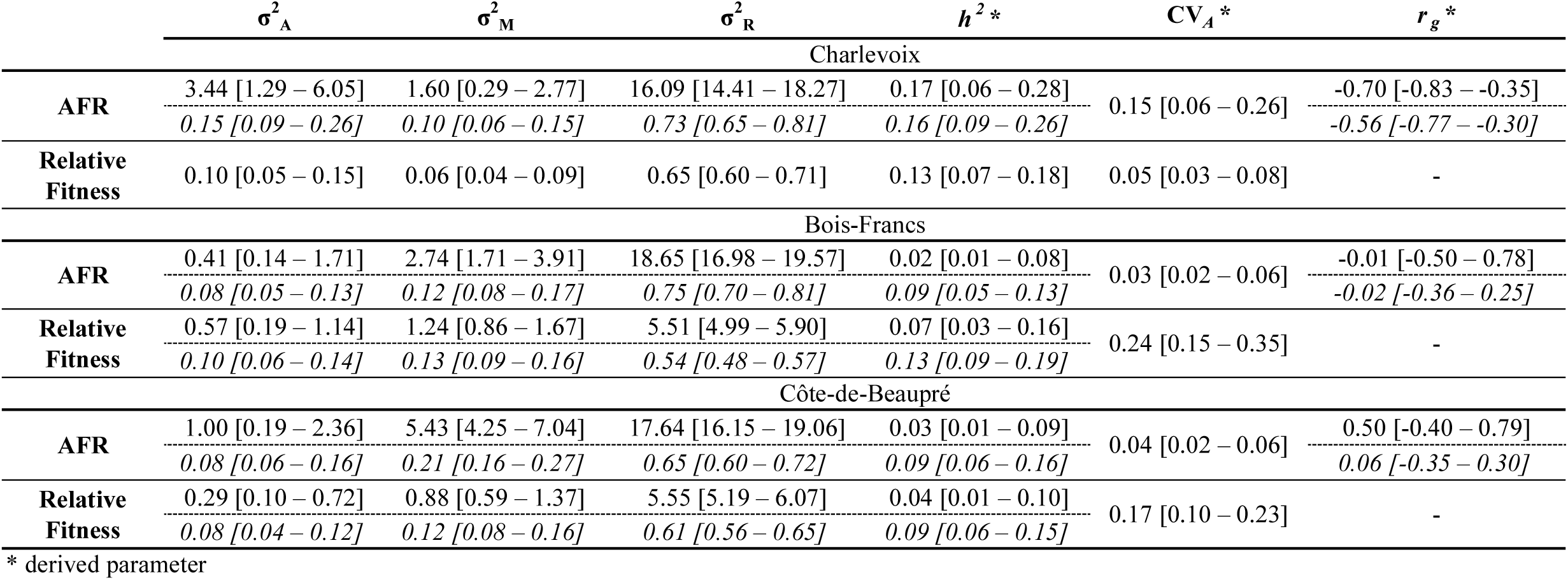
Results of the bivariate animal model fitting AFR and fitness as response variables for each dataset. **σ**^**2**^_**A**_: additive genetic variance; **σ**^**2**^_**M**_: shared familial environment variance; **σ**^**2**^_**R**_: residual variance; ***h***^***2***^: narrow-sense heritability; ***r***_***g***_: genetic correlation between AFR and relative fitness; **CV**_***A***_: coefficient of additive variation; Values are posterior modes for each parameter, along with 95% highest posterior density (HPD) intervals in brackets. Values in italic represent model results using standardized variables.

**Figure 2.**
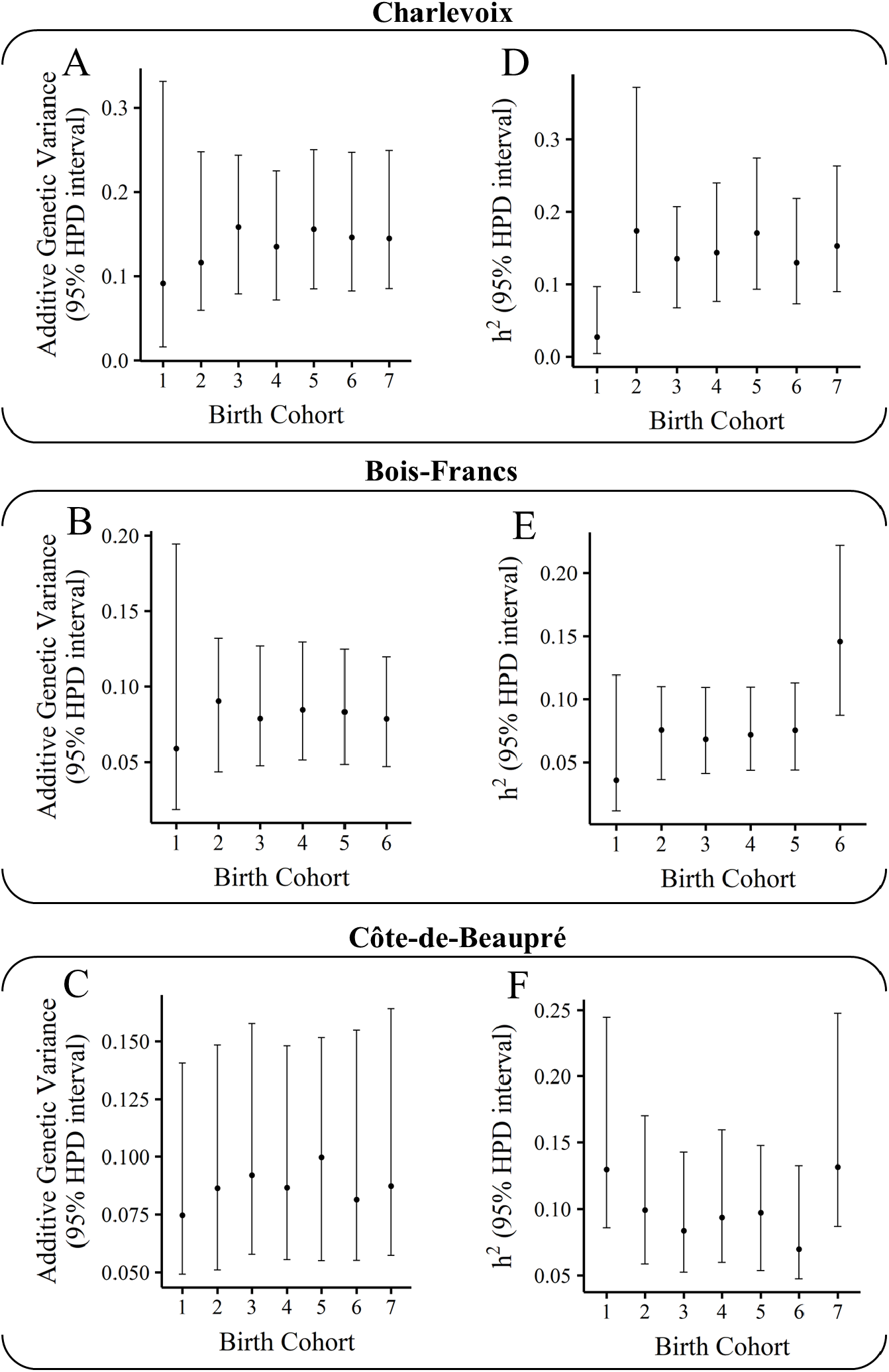
No clear pattern of temporal change in additive genetic variance of AFR, but its heritability does increase at the end of the study period. The additive genetic variance per cohort is calculated as the per-cohort variance of breeding values. Dots represent the posterior mode of the per-cohort variance with the 95% HPD interval.

The genetic correlation between AFR and relative fitness was completely different between the three populations: in Charlevoix, the genetic correlation was negative and strong in magnitude (*r*_*g*_ = −0.56 [−0.77 – −0.30]); in Bois-Francs, the correlation was negligeable (*r*_*g*_ = −0.02 [−0.36 – 0.25]), the same result was found for Côte-de-Beaupré (*r*_*g*_ = 0.06 [−0.35 to 0.30]). We found that the additive genetic covariance between AFR and fitness, which also corresponds to the STS, differed between Charlevoix and the two other populations (Table 3). For the Charlevoix population, σ_A_ = −0.07 [−0.13 – −0.02] with a *pd* index of 100%, which means that is highly likely that it is negative in direction. The ROPE (25%) index reveals that there is a *P* = 0.007 that this parameter falls in the range [−0.03 – 0.03]. For the Bois-Francs population, σ_A_ = −0.002 [−0.034 – 0.025] with a *pd* index of 59.3%, which means that is equally likely that this parameter could be either negative or positive in direction. The ROPE (25%) index indicates that there is a *P* = 0.936 that this parameter falls in the range [−0.03 – 0.03]. The same applied to the Côte-de-Beaupré population, σ_A_ = 0.007 [−0.037 – 0.026] with a *pd* index of 50.8% with a *P* = 0.939 that this parameter falls in the range [−0.03 – 0.03] (Table 4).

**Table 3.** Continued results of the bivariate animal model fitting AFR and relative fitness as response variables for each dataset. **σ**_**A**_: additive genetic covariance between AFR and relative fitness; ***β***_**G**_: genetic selection gradient; ***β***_**E**_: environmental selection gradient; ***Δβ***: bias metric in genetic selection gradient; Values are posterior modes for each parameter, along with 95% highest posterior density (HPD) intervals in brackets. Values in italic represent model results using standardized variables.

| $\sigma_A$ | $\beta_G^*$ | $\beta_E^*$ | $\Delta\beta^*$ |
| --- | --- | --- | --- |
| Charlevoix |  |  |  |
| -0.30 [-0.64 – -0.06] | -0.09 [-0.16 – -0.05] | -0.12 [-0.13 – -0.11] | -0.03 [-0.08 – 0.04] |
| *-0.07 [-0.13 – -0.02]* | *-0.37 [-0.65 – -0.22]* | *-0.55 [-0.61 – -0.51]* | *-0.19 [-0.39 – 0.11]* |
| Bois-Francis |  |  |  |
| -0.02 [-0.35 – 0.64] | -0.01 [-0.60 – 1.01] | -0.16 [-0.19 – -0.13] | -0.18 [-1.19 – 0.44] |
| *-0.002 [-0.034 – 0.025]* | *-0.03 [-0.38 – 0.32]* | *-0.24 [-0.29 – -0.20]* | *-0.18 [-0.59 – 0.17]* |
| Côte-de-Beaupré |  |  |  |
| 0.18 [-0.29 – 0.60] | 0.13 [-0.25 – 0.75] | -0.21 [-0.25 – -0.19] | -0.42 [-0.90 – 0.06] |
| *0.007 [-0.037 – 0.026]* | *0.06 [-0.34 – 0.27]* | *-0.37 [-0.42 – -0.34]* | *-0.45 [-0.69 – -0.04]* |

\* derived parameter

**Table 4.** Bayesian posterior indices for additive genetic covariance between AFR and relative fitness ***σ***_***A***_; genetic selection gradient ***β***_***G***_; environmental selection gradient ***β***_***E***_; bias metric in genetic selection gradient **Δ*β***. *pd* refers to the probability of direction, expressed here in percentage; *ROPE* refers to the proportion of probability that the 95% HPD interval that lies in the defined *Region of Practical Equivalence* based on two thresholds (10% and 25%; see Supplementary Material for details).

|  | *pd* | *ROPE* (25%) | *ROPE* (10%) |
| --- | --- | --- | --- |
| Parameter | Charlevoix |  |  |
| $\sigma_A$ | 100% | 0.007 | 0.000 |
| $\beta_G$ | 100% | 0.000 | 0.000 |
| $\beta_E$ | 100% | 0.000 | 0.000 |
| $\Delta\beta$ | 87.80% | 0.084 | 0.041 |
|  | Bois-Francis |  |  |
| - | 59.30% | 0.936 | 0.482 |
| - | 59.30% | 0.101 | 0.040 |
| - | 100% | 0.000 | 0.000 |
| - | 84.90% | 0.077 | 0.030 |
|  | Côte-de-Beaupré |  |  |
| - | 50.80% | 0.939 | 0.515 |
| - | 50.80% | 0.132 | 0.048 |
| - | 100% | 0.000 | 0.000 |
| - | 97.80% | 0.007 | 0.000 |

We used the approach of calculating the metric of bias in the standardized phenotypic selection gradient to reveal if there is any environmental bias. For the population of Charlevoix, we found *Δβ* = −0.19 [−0.39 – 0.11] (Table 3). According to the ROPE (25%) index, there is a *P* = 0.084 that *Δβ* of Charlevoix would fall in the range [−0.03 – 0.03]. For the Bois-Francs population, *Δβ* = −0.18 [−0.59 – 0.17] with a *P* = 0.077 that it would fall in the range [−0.03 – 0.03]. For the Côte-de-Beaupré population, *Δβ* = −0.45 [−0.69 – −0.04] with a *P* = 0.007 that it would fall in the range [−0.03 – 0.03] (Table 4).

### Observed vs predicted microevolutionary change

The results from the animal models clearly reveal that AFR is heritable (Figure 2), under unbiased selection and strongly correlated with our fitness measure. Using STS, given by the additive genetic covariance between AFR and fitness, σ_A_, we found that the expected microevolutionary change from one generation to the next in AFR is an advancement of 0.07 phenotypic standard deviations in Charlevoix (or the equivalent of 25.5 days/per generation, or ½ a year for the whole study period), an advancement of 0.002 phenotypic standard deviations in Bois-Francs (or the equivalent of 0.73 days/per generation, or the equivalent of 4 days for the whole study period), and an increase of 0.007 phenotypic standard deviations in Côte-de-Beaupré (or the equivalent of 2.5 days/per generation, or the equivalent of 17.5 days for the whole study period). Using BE, given by multiplying the standardized phenotypic covariance between AFR and fitness by the trait’s heritability, we found an expected advancement in AFR of 0.09 standard deviations in Charlevoix (or the equivalent 34.65 days/per generation, or the equivalent of ⅔ a year for the whole study period), an advancement of around 0.03 phenotypic standard deviations in both Bois-Francs and Côte-de-Beaupré (or the equivalent of around 10.5 days/per generation, or the equivalent of 63 and 73.5 days for the whole study period, respectively). The prediction based on the breeder’s equation is ∼50% higher than those made with the STS.

To establish if there was a trend in the average AFR, we ran a mixed effects linear regression model with birth cohort fitted as a covariate. Results from these models show that all three populations had completely different temporal trends in average AFR (Figures 1 and 3). The partial regression coefficient of AFR on birth cohort (i.e., slope of regression line) in Charlevoix, *β* = 0.02 [−0.01 – 0.06], in Bois-Francs, *β* = −0.17 [−0.20 – −0.13], and in Côte-de-Beaupré, *β* = 0.08 [0.06 – 0.10]. We examined the trends in the average breeding values of AFR across the three populations to determine if there was a genetic response to selection. In all three populations, the regression coefficient of the average breeding values on birth cohort was negligible, in Charlevoix, *β*_BV_ = −0.01 [−0.04 – 0.02], in Bois-Francs, *β*_BV_ = −0.01 [−0.04 – 0.02], and in Côte-de-Beaupré, *β*_BV_ = −0.002 [−0.01 – 0.006] (Figure 3). We found that for all three populations there is a posterior probability *P* ≈ 0.5 of observing trends of these magnitude or less under a pure drift scenario.

**Figure 3.**
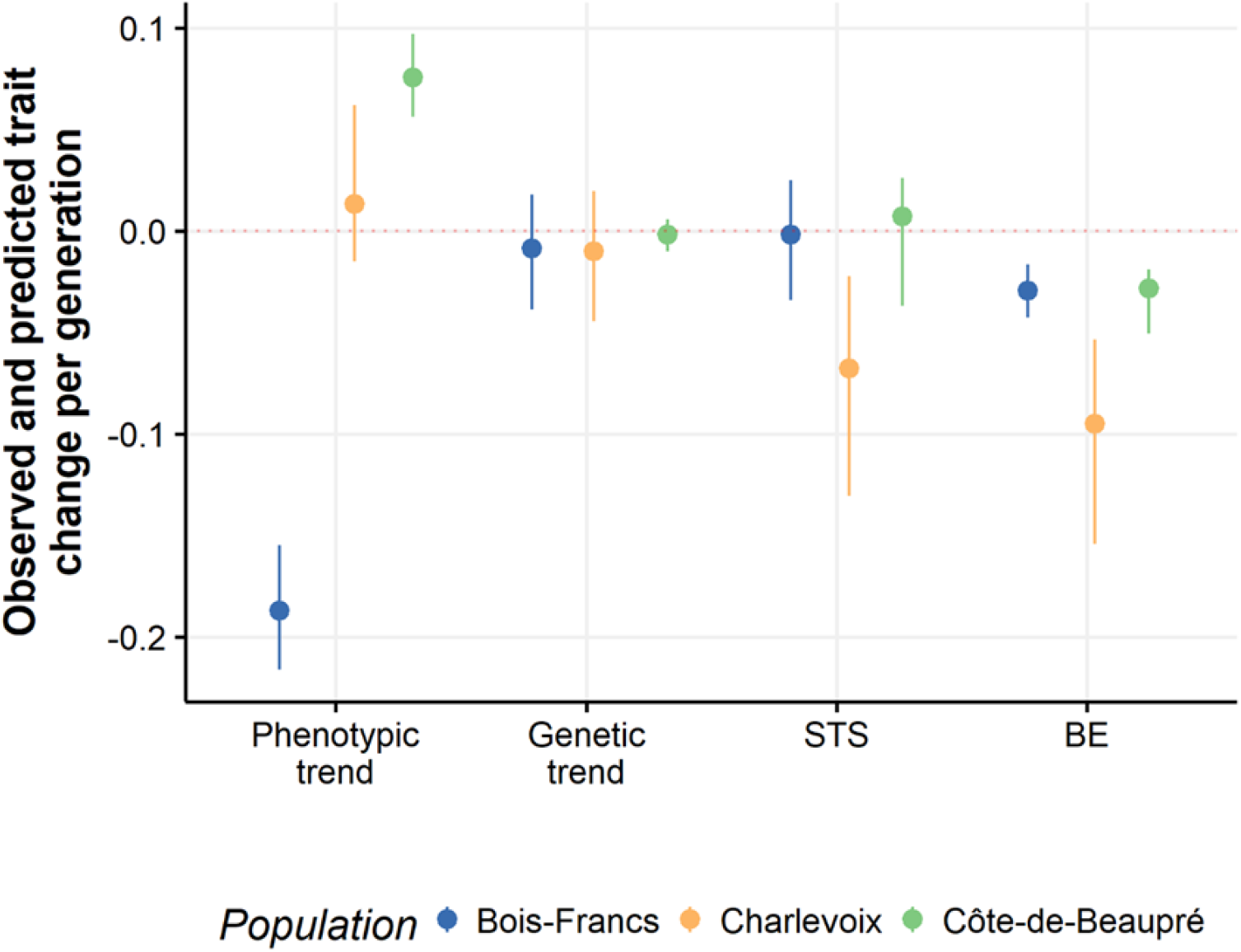
Evidence of evolutionary constrain in AFR Charlevoix and evolutionary stasis in Bois-Francs and Côte-de-Beaupré across a period of at least 150 years. The phenotypic trend represents the slope of the temporal trend in cohort. The genetic trend is based on best-fit animal model estimate and represent the slope of the temporal trend in mean breeding value of AFR. STS and BE give predicted per-generation change in mean AFR. The dots represent posterior mode-estimates, along with the 95% HPD intervals.

## Discussion

In this study, we evaluated the predictability of evolutionary outcomes in a key human life history trait, age at first reproduction (AFR), across three closely related French-Canadian populations separated at a subregional scale. Despite consistent selection favoring earlier AFR and modest heritability (*h*^2^ ≈ 0.11), we observed no uniform genetic or phenotypic response to selection. This stands in contrast to the previously studied Île aux Coudres (IAC) population, which belong to the larger Charlevoix population, where AFR declined over 140 years, accompanied by a corresponding genetic shift (Milot et al. 2011). Our findings challenge the assumption that consistent selection reliably yields evolutionary change, highlighting the uncertainty and diversity of outcomes even under seemingly parallel demographic and environmental conditions.

### Predictive Models and Evolutionary Inconsistency

Quantitative genetic theory provides a set of expectations, through the Breeder’s Equation (BE) and Robertson’s Secondary Theorem of Selection (STS), for predicting evolutionary responses under directional selection. In Charlevoix, both models predicted a decline in AFR, yet neither phenotypic means nor breeding values showed significant change. In Côte-de-Beaupré, AFR slightly increased; in Bois-Francs, it declined, but not due to genetic causes. These divergent outcomes emphasize that microevolutionary predictions, while theoretically grounded, may fail in practice, especially when genetic and environmental components interact in complex ways.

STS, by isolating the additive genetic covariance between a trait and fitness, offers a more direct test of evolutionary response than BE, which relies on total phenotypic selection. In our analyses, STS yielded more conservative predictions than BE, yet still anticipated evolutionary change that did not materialize in the Charlevoix population. This underscores a critical point: selection on heritable traits does not guarantee evolutionary response.

### What Limits Evolutionary Predictability?

The absence of a detectable genetic response in three populations suggests that AFR evolution is not only variable but also unpredictable, even across genetically and demographically similar populations. In Bois-Francs and Côte-de-Beaupré, strong selection acted on environmental deviations of AFR rather than on breeding values—a pattern consistent with cryptic evolution or evolutionary stasis (Rausher 1992; Stinchcombe et al. 2014). In Charlevoix, despite selection on both genetic and environmental components, no trait change occurred, suggesting evolutionary constraint.

These inconsistencies have important implications for demographic forecasting. Traits like AFR are key inputs in population projection models, particularly in evolutionary demography. If selection does not lead to predictable trait evolution, then demographic models assuming such change may misestimate future fertility rates, generation times, or growth trajectories.

Biologically, several mechanisms may account for these constraints. First, genotype-by-environment interactions (G×E), as previously shown in Île aux Coudres (Mawass et al. 2022), can obscure selection effects by altering trait–fitness relationships across contexts. Second, genetic correlations may constrain evolution if AFR is genetically linked to other fitness-relevant traits, creating trade-offs (Arnold et al. 2008). Third, demographic factors, such as inbreeding and small effective population sizes (*N*_*e*_), may reduce the available additive genetic variance or increase the influence of random genetic drift.

Despite these complexities, our animal model analyses confirm that AFR retains evolutionary potential: heritability was detectable, and evolvability measures suggested sufficient genetic variance for change—yet no response occurred. This highlights a core issue in evolutionary biology: quantitative genetic potential does not always translate into evolutionary reality, especially in natural populations where selective pressures, environmental variance, and genetic architecture interact in nonlinear ways.

### Methodological Robustness and Limitations

We took steps to ensure the robustness of our conclusions. Sensitivity analyses confirmed that our pedigrees had sufficient power to detect genetic parameters and low genetic correlations (*r*_*g*_ ≥ 0.1), minimizing the likelihood that failure to detect evolution reflects statistical error (Mawass and Milot 2025). While Bayesian priors help constrain uncertainty, they may introduce bias; however, the consistency of our results across models and populations lends credibility to our conclusions. Still, the interpretation of null evolutionary responses requires caution, especially when statistical and biological sources of constraint are intertwined (Pujol et al. 2018).

### Conclusion: Predictability, Not Generality, of Evolution

Our findings suggest that evolutionary responses to selection on key life history traits are not reliably predictable, even among closely related populations facing similar environments. This has fundamental implications for how we understand and model evolution in human populations. Rather than assuming that microevolutionary change is a consistent outcome of selection, we propose that evolutionary stasis or divergence should be the default expectation, unless demonstrated otherwise.

In applied contexts—such as evolutionary demography, public health, or life history modeling—this uncertainty must be accounted for. Predictive models of human evolution that ignore sources of evolutionary constraint risk misestimating future population dynamics. As evolutionary biology increasingly moves toward forecasting frameworks (Hendry et al. 2011; Kingsolver and Diamond 2011), recognizing the limits of predictability will be essential for developing robust, realistic models.

Future research should combine multivariate quantitative genetic approaches with genetic data and environmental covariates to identify the causes of constrained evolution. Integrating G-matrix structures, G×E effects, and fitness trade-offs will improve our understanding of why selection does not always result in genetic change. In doing so, we can move from describing evolutionary outcomes to explaining—and eventually predicting—them with greater precision.

## Author Contributions

WM conceived and developed the study; performed data and computational analysis with input from EM; and drafted the manuscript. EM contributed to the final version of the manuscript. EM supervised and funded the project.

## Acknowledgements

We acknowledge the BALSAC Project at the Université du Québec à Chicoutimi for maintaining and supporting its demographic database. The study would not have been possible without the foundational work of Gérard Bouchard, who initiated the BALSAC Project in 1976 and established its population registry. These organizations played no role in the design, analysis, interpretation, or reporting of this research.

## Data Availability

The data are part of the BALSAC project that contains vital information. Due to constraints related to regulations of access to personal information, the pedigree data are accessible only upon request.

## References

Anderson-Trocmé, Luke, Dominic Nelson, Shadi Zabad, et al. 2023. “On the Genes, Genealogies, and Geographies of Quebec.” Science 380 (6647): 849–55. 10.1126/science.add5300.

Arnold, Stevan J., Reinhard Bürger, Paul A. Hohenlohe, et al. 2008. “Understanding the Evolution and Stability of the G-Matrix.” Evolution 62 (10): 10. 10.1126/scisignal.2001449.Engineering.

Barrett, Rowan D. H., Antoine Paccard, Timothy M. Healy, et al. 2010. “Rapid Evolution of Cold Tolerance in Stickleback.” Proceedings of the Royal Society B: Biological Sciences 278 (1703): 233–38. 10.1098/rspb.2010.0923.

Barton, John P., Nilu Goonetilleke, Thomas C. Butler, Bruce D. Walker, Andrew J. McMichael, and Arup K. Chakraborty. 2016. “Relative Rate and Location of Intra-Host HIV Evolution to Evade Cellular Immunity Are Predictable.” Nature Communications 7 (1): 11660. 10.1038/ncomms11660.

Boisvert, Mireille, and Francine M. Mayer. 1994. “Mortalité Infantile et Consanguinité Dans Une Population Endogame Du Québec.” Population 49 (3): 3.

Bolund, Elisabeth, Adam Hayward, Jenni E. Pettay, and Virpi Lummaa. 2015. “Effects of the Demographic Transition on the Genetic Variances and Covariances of Human Life-History Traits.” Evolution 69 (3): 747–55. 10.1111/evo.12598.

Bonnet, Timothée, Michael B. Morrissey, Alison Morris, et al. 2019. “The Role of Selection and Evolution in Changing Parturition Date in a Red Deer Population.” PLOS Biology 17 (11): e3000493. 10.1371/journal.pbio.3000493.

Bonnet, Timothée, Michael B. Morrissey, Pierre de Villemereuil, et al. 2022. “Genetic Variance in Fitness Indicates Rapid Contemporary Adaptive Evolution in Wild Animals.” Science 376 (6596): 1012–16. 10.1126/science.abk0853.

Bouchard, Gérard, Marc de Braekeleer, and SOREP. 1991. Histoire d’un génôme: population et génétique dans l’est du Québec. Presses de l’Université du Québec.

Bouchard, Grard, Raymond Roy, Bernard Casgrain, and Michel Hubert. 1989. “Fichier de Population et Structures de Gestion de Base de Données: Le Fichier-Réseau BALSAC et Le Système INGRES/INGRID.” Histoire & Mesure 4 (1/2): 39–57.

Byars, Sean G., Douglas Ewbank, Diddahally R. Govindaraju, and Stephen C. Stearns. 2010. “Colloquium Papers: Natural Selection in a Contemporary Human Population.” Proceedings of the National Academy of Sciences 107 Suppl (October): October. 10.1073/pnas.0906199106.

Byars, Sean G., and Konstantinos Voskarides. 2019. “Genes That Improved Fitness Also Cost Modern Humans: Evidence for Genes with Antagonistic Effects on Longevity and Disease.” Evolution, Medicine and Public Health 2019 (1): 1. 10.1093/emph/eoz002.

Colautti, Robert I., and Jennifer A. Lau. 2015. “Contemporary Evolution during Invasion: Evidence for Differentiation, Natural Selection, and Local Adaptation.” Molecular Ecology 24 (9): 1999–2017. 10.1111/mec.13162.

Cole, Lamont C. 1954. “The Population Consequences of Life History Phenomena.” The Quarterly Review of Biology 29 (2): 103–37. 10.1086/400074.

De Villemereuil, Pierre, Holger Schielzeth, Shinichi Nakagawa, and Michael B. Morrissey. 2016. “General Methods for Evolutionary Quantitative Genetic Inference from Generalized Mixed Models.” Genetics 204 (3): 3. 10.1534/genetics.115.186536.

Dillon, Lisa, Marilyn Amorevieta-Gentil, Marianne Caron, et al. 2018. “The Programme de Recherche En Démographie Historique: Past, Present and Future Developments in Family Reconstitution.” The History of the Family 23 (1): 20–53. 10.1080/1081602X.2016.1222501.

Engelhardt, Sacha C., Patrick Bergeron, Alain Gagnon, Lisa Dillon, and Fanie Pelletier. 2019. “Using Geographic Distance as a Potential Proxy for Help in the Assessment of the Grandmother Hypothesis.” Current Biology 29 (4): 4. 10.1016/j.cub.2019.01.027.

Fisher, Ronald A. 1930. The Genetical Theory of Natural Selection. Vol. 154. Clarendon. http://openlibrary.org/books/OL7084333M.

Gold, Ellen B. 2011. “The Timing of the Age at Which Natural Menopause Occurs.” Obstetrics and Gynecology Clinics of North America 38 (3): 425–40. 10.1016/j.ogc.2011.05.002.

Grant, Peter R., and B. Rosemary Grant. 2002. “Unpredictable Evolution in a 30-Year Study of Darwin’s Finches.” Science 296 (5568): 707–11. 10.1126/science.1070315.

Hadfield, Jarrod D. 2010. “MCMC Methods for Multi-Respoinse Generalized Linear Mixed Models: The MCMCglmm R Package.” Journal of Statistical Software 33 (2): 2. 10.1002/ana.22635.

Hadfield, Jarrod D, Alastair J Wilson, and Loeske E B Kruuk. 2011. “Cryptic Evolution: Does Environmental Deterioration Have a Genetic Basis?” Genetics 187 (4): 1099–113. 10.1534/genetics.110.124990.

Hansen, Thomas F., and Houle David. 2004. “Evolvability, Stabilizing Selection, and the Problem of Stasis.” Phenotypic Integration, April, 130–52. 10.1093/oso/9780195160437.003.0006.

Henderson, C. R. 1975. “Best Linear Unbiased Estimation and Prediction under a Selection Model.” Biometrics 31 (2): 423–47. 10.2307/2529430.

Henderson, C. R. 1986. “Estimation of Variances in Animal Model and Reduced Animal Model for Single Traits and Single Records.” Journal of Dairy Science 69 (5): 1394–402. 10.3168/jds.S0022-0302(86)80546-X.

Hendry, Andrew P., and Michael T. Kinnison. 1999. “Perspective: The Pace of Modern Life : Measuring Rates of Contemporary Microevolution.” Evolution 53 (6): 6.

Hendry, Andrew P., Michael T. Kinnison, Mikko Heino, et al. 2011. “Evolutionary Principles and Their Practical Application.” Evolutionary Applications 4 (2): 159–83. 10.1111/j.1752-4571.2010.00165.x.

Hendry, Andrew P., Daniel J. Schoen, Matthew E. Wolak, and Jane M. Reid. 2018. “The Contemporary Evolution of Fitness.” Annual Review of Ecology Evolution and Systematics 49: 457–76. 10.1146/annurev-ecolsys-110617.

Houle, David. 1992. “Comparing Evolvability and Variability of Quantitative Traits.” Genetics 130: 195–204.

Kingsolver, Joel G., and Sarah E. Diamond. 2011. “Phenotypic Selection in Natural Populations: What Limits Directional Selection?” The American Naturalist 177 (3): 346–57. 10.1086/658341.

Kosova, G., M. Abney, and C. Ober. 2010. “Heritability of Reproductive Fitness Traits in a Human Population.” Proceedings of the National Academy of Sciences 107 (Suppl_1): suppl_1. 10.1073/pnas.0906196106.

Kruschke, John. 2014. Doing Bayesian Data Analysis: A Tutorial with R, JAGS, and Stan. Academic Press.

Kruuk, Loeske E. B. 2004. “Estimating Genetic Parameters in Natural Populations Using the ‘Animal Model.’” Philosophical Transactions of the Royal Society of London B Biological Sciences 359 (1446): 1446. 10.1098/rstb.2003.1437.

Kruuk, Loeske E. B., Julianne Livingston, Andrew Kahn, and Michael D. Jennions. 2015. “Sex-Specific Maternal Effects in a Viviparous Fish.” Biology Letters 11 (8): 20150472. 10.1098/rsbl.2015.0472.

Lande, Russell, and Stevan J. Arnold. 1983. “The Measurement of Selection on Correlated Characters.” Evolution 37 (6): 6. 10.2307/2408842.

Lässig, Michael, Ville Mustonen, and Aleksandra M. Walczak. 2017. “Predicting Evolution.” Nature Ecology & Evolution 1 (3): 0077. 10.1038/s41559-017-0077.

Lenski, Richard E., and Philip M. Service. 1982. “The Statistical Analysis of Population Growth Rates Calculated from Schedules of Survivorship and Fecunidity.” Ecology 63 (3): 655– 62. 10.2307/1936785.

Lush, Jay Laurence. 1937. Animal Breeding Plans.

Lynch, Michael, and Bruce Walsh. 1998. Genetics and Analysis of Quantitative Traits. Sinauer.

Makowski, Dominique, Mattan S. Ben-Shachar, S. H.Annabel Chen, and Daniel Lüdecke. 2019. “Indices of Effect Existence and Significance in the Bayesian Framework.” Frontiers in Psychology 10 (December): December. 10.3389/fpsyg.2019.02767.

Makowski, Dominique, Mattan S. Ben-Shachar, and Daniel Lüdecke. 2019. “bayestestR: Describing Effects and Their Uncertainty, Existence and Significance within the Bayesian Framework.” Journal of Open Source Software 4 (40): 1541. 10.21105/joss.01541.

Mawass, Walid, Francine M. Mayer, and Emmanuel Milot. 2022. “Genotype-by-Environment Interactions Modulate the Rate of Microevolution in Reproductive Timing in Humans.” Evolution 76 (7): 1391–405. 10.1111/evo.14504.

Mawass, Walid, and Emmanuel Milot. 2025. “Assessing the Impact of Pedigree Attributes on the Validity of Quantitative Genetic Parameter Estimates.” Journal of Evolutionary Biology 38 (4): 439–56. 10.1093/jeb/voaf010.

Milot, Emmanuel, Francine M. Mayer, Daniel H. Nussey, Mireille Boisvert, Fanie Pelletier, and Denis Réale. 2011. “Evidence for Evolution in Response to Natural Selection in a Contemporary Human Population.” Proceedings of the National Academy of Sciences 108 (41): 41. 10.1073/pnas.1104210108.

Moorad, Jacob A. 2013. “A Demographic Transition Altered the Strength of Selection for Fitness and Age-Specific Survival and Fertility in a 19th Century American Population.” Evolution 67 (6): 6. 10.1111/evo.12023.

Moorad, Jacob A. 2014. “Individual Fitness and Phenotypic Selection in Age-Structured Populations with Constant Growth Rates.” Ecology 95 (4): 4. 10.1890/13-0778.1.

Moreau, Claudia, Claude Bhérer, Hélène Vézina, Michèle Jomphe, Damian Labuda, and Laurent Excoffier. 2011. “Deep Human Genealogies Reveal a Selective Advantage to Be on an Expanding Wave Front.” Science 334 (6059): 6059. 10.1126/science.1212880.

Morrissey, Michael B., and Jarrod D. Hadfield. 2012. “Directional Selection in Temporally Replicated Studies Is Remarkably Consistent.” Evolution 66 (2): 2. 10.1111/j.1558-5646.2011.01444.x.

Morrissey, Michael B., Darren J. Parker, Peter Korsten, Josephine M. Pemberton, Loeske E. B. Kruuk, and Alastair J. Wilson. 2012. “The Prediction of Adaptive Evolution: Empirical Application of the Secondary Theorem of Selection and Comparison to the Breeder’s Equation.” Evolution 66 (8): 8. 10.1111/j.1558-5646.2012.01632.x.

Neher, Richard A, Colin A Russell, and Boris I Shraiman. 2014. “Predicting Evolution from the Shape of Genealogical Trees.” eLife 3 (November): e03568. 10.7554/eLife.03568.

Nosil, Patrik, Romain Villoutreix, Clarissa F. De Carvalho, et al. 2018. “Natural Selection and the Predictability of Evolution in Timema Stick Insects.” Science 359 (6377): 765–70. 10.1126/science.aap9125.

Nussey, Daniel H., Alastair J. Wilson, and Jon E. Brommer. 2007. “The Evolutionary Ecology of Individual Phenotypic Plasticity in Wild Populations.” Journal of Evolutionary Biology 20 (3): 3. 10.1111/j.1420-9101.2007.01300.x.

Otto, Sarah P. 2018. “Adaptation, Speciation and Extinction in the Anthropocene.” Proceedings of the Royal Society B: Biological Sciences 285 (1891): 1891. 10.1098/rspb.2018.2047.

Pavard, Samuel, and Christophe F. D. Coste. 2021. “Evolutionary Demographic Models Reveal the Strength of Purifying Selection on Susceptibility Alleles to Late-Onset Diseases.” Nature Ecology & Evolution 5 (3): 392–400. 10.1038/s41559-020-01355-2.

Peischl, Stephan, Isabelle Dupanloup, Adrien Foucal, et al. 2018. “Relaxed Selection During a Recent Human Expansion.” Genetics 208 (2): 763–77. 10.1534/genetics.117.300551.

Pelletier, Fanie, Gabriel Pigeon, Patrick Bergeron, et al. 2017. “Eco-Evolutionary Dynamics in a Contemporary Human Population.” Nature Communications 8 (May): May. 10.1038/ncomms15947.

Pigeon, Gabriel, Marco Festa-Bianchet, David W. Coltman, and Fanie Pelletier. 2016. “Intense Selective Hunting Leads to Artificial Evolution in Horn Size.” Evolutionary Applications 9 (4): 521–30. 10.1111/eva.12358.

Price, Trevor, Mark Kirkpatrick, and Steven J. Arnold. 1988. “Directional Selection and the Evolution of Breeding Date in Birds.” Science 240 (4853): 798–99. 10.1126/science.3363360.

Pujol, Benoit, Simon Blanchet, Anne Charmantier, et al. 2018. “The Missing Response to Selection in the Wild.” Trends in Ecology & Evolution 33 (5): 337–46. 10.1016/j.tree.2018.02.007.

Rausher, Mark D. 1992. “The Measurement of Selection on Quantitative Traits: Biases Due to Environmental Covariances between Traits and Fitness.” Evolution 46 (3): 616–26. 10.1111/j.1558-5646.1992.tb02070.x.

Robertson, Alan. 1966. “A Mathematical Model of the Culling Process in Dairy Cattle.” Animal Science 8 (1): 95–108. 10.1017/S0003356100037752.

Robertson, Alan. 1968. “The Spectrum of Genetic Variation.” Population Biology and Evolution, 5–16.

Roff, Derek. 1993. Evolution Of Life Histories: Theory and Analysis. Springer Science & Business Media.

Sanjak, Jaleal S., Julia Sidorenko, Matthew R. Robinson, Kevin R. Thornton, and Peter M. Visscher. 2017. “Evidence of Directional and Stabilizing Selection in Contemporary Humans.” Proceedings of the National Academy of Sciences 115 (1): 1. 10.1073/pnas.1707227114.

Sauve, Drew, George Divoky, and Vicki L. Friesen. 2019. “Phenotypic Plasticity or Evolutionary Change? An Examination of the Phenological Response of an Arctic Seabird to Climate Change.” Functional Ecology 33 (11): 2180–90. 10.1111/1365-2435.13406.

Siepielski, Adam M., Joseph D. DiBattista, and Stephanie M. Carlson. 2009. “It’s about Time: The Temporal Dynamics of Phenotypic Selection in the Wild.” Ecology Letters 12 (11): 1261– 76. 10.1111/j.1461-0248.2009.01381.x.

Siepielski, Adam M., Kiyoko M. Gotanda, Michael B. Morrissey, Sarah E. Diamond, Joseph D. DiBattista, and Stephanie M. Carlson. 2013. “The Spatial Patterns of Directional Phenotypic Selection.” Ecology Letters 16 (11): 1382–92. 10.1111/ele.12174.

Stearns, Stephen C. 1992. The Evolution of Life Histories. Oxford University Press.

Stern, Aaron J., Leo Speidel, Noah A. Zaitlen, and Rasmus Nielsen. 2021. “Disentangling Selection on Genetically Correlated Polygenic Traits via Whole-Genome Genealogies.” The American Journal of Human Genetics 108 (2): 219–39. 10.1016/j.ajhg.2020.12.005.

Stinchcombe, John R., Anna K. Simonsen, and Mark. W. Blows. 2014. “Estimating Uncertainty in Multivariate Responses to Selection.” Evolution 68 (4): 1188–96. 10.1111/evo.12321.

Stuart, Y. E., T. S. Campbell, P. A. Hohenlohe, R. G. Reynolds, L. J. Revell, and J. B. Losos. 2014. “Rapid Evolution of a Native Species Following Invasion by a Congener.” Science 346 (6208): 463–66. 10.1126/science.1257008.

Teplitsky, Celine, Maja Tarka, Anders >P. Møller, et al. 2014. “Assessing Multivariate Constraints to Evolution across Ten Long-Term Avian Studies.” PLOS ONE 9 (3): e90444. 10.1371/journal.pone.0090444.

Tropf, Felix C., Gert Stulp, Nicola Barban, et al. 2015. “Human Fertility, Molecular Genetics, and Natural Selection in Modern Societies.” PLoS ONE 10 (6): 6. 10.1371/journal.pone.0126821.

Vézina, Hélène, and Jean-Sébastien Bournival. 2020. “An Overview of the BALSAC Population Database. Past Developments, Current State and Future Prospects.” Historical Life Course Studies 9 (August): 114–29. 10.51964/hlcs9299.

Wilson, Alastair J., Denis Réale, Michelle N. Clements, et al. 2010. “An Ecologist’s Guide to the Animal Model.” Journal of Animal Ecology 79 (1): 1. 10.1111/j.1365-2656.2009.01639.x.

Wu, Yihan, and Robert I. Colautti. 2022. “Evidence for Continent-Wide Convergent Evolution and Stasis throughout 150 y of a Biological Invasion.” Proceedings of the National Academy of Sciences 119 (18): e2107584119. 10.1073/pnas.2107584119.

