## Supplemental Information for "Contrasting evolutionary outcomes in a human life history trait which is heritable and under consistent unbiased directional selection"

### Testing the "migration" hypothesis

Civil acts do not allow to distinguish emigrants, who spent their reproductive life partly or in totality in the area, from less or non-fecund couples (Boisvert and Mayer 1994). To tackle this problem, we applied the data filtering previously used in other studies on a similar population (île aux Coudres; Boisvert and Mayer, 1994; Milot et al. 2011; Pelletier et al. 2017). Using this data filtering, we generated two different datasets for each population based on different assumptions regarding the unusually long interbirth intervals in the records. The "subfecundity" dataset assumes that unusually long interbirth intervals reflect subfecundity. The "migration" dataset assumes that long intervals may also reflect emigration from the region and excludes families with such length intervals. Since we detected no noticeable differences between the results obtained under these two hypotheses in all three populations, we present only the results for the subfecundity hypothesis.

### Highest posterior density (HPD) interval

The 95% highest posterior density (HPD) interval represents the most compact range of parameter values encompassing 95% of the total posterior probability mass. It is important to clarify that this interval does not uniquely identify the region most likely to contain the true parameter value, any interval covering 95% of the posterior (such as one beginning at the lower tail and accumulating upward) has the same probability of enclosing the true value. What distinguishes the HPD interval is that, under a unimodal posterior, all values within it have higher posterior density than those outside. When the posterior mode is used as the central estimate, the HPD offers a concise summary of the associated uncertainty. However, HPD intervals should not be confused with 95% confidence intervals from frequentist inference. In particular, the inclusion or exclusion of zero in an HPD interval does not imply "statistical significance" in the classical sense. Instead, complementary Bayesian measures such as the probability of direction (pd) provide a more appropriate bridge between Bayesian estimates and their frequentist interpretations, which we have employed in our work.

**Assigning the Region of Practical Equivalence (*ROPE*)**

The Region of Practical Equivalence (ROPE) quantifies how much of the HPD interval falls within a predefined range of parameter values that are considered too small to be biologically meaningful (as discussed in Makowski et al. 2019b). For example, suppose we define genetic correlations between –0.1 and 0.1 as biologically negligible. If the 95% HPD interval for a genetic correlation estimate spans from 0.02 to 0.3, but only 80% of that distribution lies outside the ROPE (i.e., beyond |0.1|), then the probability that the true correlation exceeds the biologically negligible range is just 0.8. Although this interval excludes zero, something that might be interpreted as "significant" under frequentist thinking, the ROPE framework provides a more nuanced Bayesian interpretation of practical importance.

In this study, we employed the ROPE index to evaluate the magnitude of parameters from our animal models, interpreting them in terms of their biological relevance. While the choice of ROPE boundaries is ultimately subjective, we transparently justified our approach. Recognizing that different researchers may have different views on what constitutes a meaningful effect size, we applied three distinct thresholds: liberal, moderate, and conservative. The ROPE thresholds were applied to additive genetic covariance components, genetic selection gradient, environmental selection gradient, and bias metric in genetic selection gradient. Specifically, we used: (1) a liberal threshold of 5% of the total estimate (e.g., assuming $\boldsymbol{\sigma_A}$ of 0.1, any value ≥0.005 would be considered meaningful), (2) a conservative threshold of 25%, requiring a larger component contribution to be deemed non-negligible, and (3) a moderate cutoff of 10%, providing an intermediate standard of practical importance. In the main text, we present the results of the 25% and 10% cutoffs (Table 4).

### Tables

**Table S1**. Results from the phenotypic mixed effect linear regression model of fitness as a function of AFR, for each tested population. Values are posterior modes for each parameter, along with 95% highest posterior density (HPD) intervals in brackets. All variables in this analysis are standardized to allow for among population comparisons.

| | **Population** | | |
|---|---|---|---|
| | Charlevoix | Bois-Francs | Côte-de-Beaupré |
| **Fixed effect** | | | |
| Intercept | -0.98 [-1.15 – -0.78] | -1.54 [-1.72 – -1.38] | -0.88 [-1.00 – -0.67] |
| AFR (linear) | -0.64 [-0.68 – -0.60] | -0.25 [-0.29 – -0.22] | -0.34 [-0.37 – -0.30] |
| AFR (quadratic) | 0.07 [0.05 – 0.08] | 0.02 [0.003 – 0.04] | 0.002 [-0.02 – 0.02] |
| Inbreeding (linear) | 0.03 [0.05 – 0.08] | -0.02 [-0.07 – 0.02] | 0.17 [0.12 – 0.22] |
| Inbreeding (quadratic) | -0.008 [-0.02 – 0.008] | 0.001 [-0.003 – 0.008] | -0.02 [-0.04 – -0.01] |
| Birth cohort | 0.14 [0.12 – 0.17] | 0.35 [0.32 – 0.38] | 0.23 [0.21 – 0.24] |
| Twin birth | 0.42 [0.31 – 0.50] | 0.45 [0.34 – 0.53] | 0.35 [0.24 – 0.43] |
| Infant mortality | 0.27 [0.25 – 0.31] | 0.06 [0.03 – 0.08] | 0.16 [0.13 – 0.19] |
| Parish | -0.03 [-0.14 – 0.08] | -0.44 [-0.86 – 0.02] | 0.40 [-0.16 – 0.91] |
| **Variance component** | | | |
| Maternal environment | 0.04 [0.002 – 0.06] | 0.14 [0.10 – 0.16] | 0.10 [0.06 – 0.13] |
| Residual | 0.45 [0.42 – 0.49] | 0.54 [0.50 – 0.57] | 0.59 [0.54 – 0.62] |
